# Dietary calcium (Ca^2+^) impacts Ca^2+^ content and molecular expression of Ca^2+^-transporters in Malpighian tubules of the yellow fever mosquito, *Aedes aegypti*

**DOI:** 10.1101/2023.03.30.534971

**Authors:** Yuan Li, Peter M. Piermarini

## Abstract

The renal (Malpighian) tubules of insects play important roles in hemolymph Ca^2+^ regulation. Here we provide the first investigation into how dietary Ca^2+^ loads from sucrose or blood meals affect the Ca^2+^ content and mRNA expression of Ca^2+^ transporters in Malpighian tubules of adult female mosquitoes. Using the yellow fever mosquito *Aedes aegypti* we found that feeding females for 6 days *ad libitum* on 10% sucrose with elevated Ca^2+^ concentration led to concentration-dependent increases of the Ca^2+^ content in Malpighian tubules. The increases of Ca^2+^ content correlated with up-regulations of mRNAs encoding intracellular Ca^2+^-ATPases (*SERCA* and *SPCA*), a plasma membrane Ca^2+^ -ATPase (*PMCA*), and a K^+^-dependent Na^+^/Ca^2+^ exchanger (*NCKX1*). We also found that when adult females were fed blood, tubule Ca^2+^ content changed dynamically over the next 72 h in a manner consistent with redistribution of tubule Ca^2+^ stores to other tissues (e.g., ovaries). The changes in tubule Ca^2+^ were correlated with dynamic changes in mRNA abundances of *SERCA*, *SPCA*, *PMCA*, and *NCKX1*. Our results are the first to demonstrate that Malpighian tubules of adult female mosquitoes have a remarkable capacity to handle high dietary Ca^2+^ loads, most likely through the combination of storing excess Ca^2+^ within intracellular compartments and secreting it into the tubule lumen for excretion. Our results also suggest that the Malpighian tubules play key roles in supplying Ca^2+^ to other tissues during the processing of blood meals.

## 1. Introduction

Renal (Malpighian) tubules of insects play key roles in maintaining hemolymph osmotic and ionic balance, as well as in detoxifying and/or excreting endogenous metabolic/nitrogenous wastes and xenobiotics (Beyenbach and Piermarini, 2009; Nocelli et al., 2016). In most insects, the quintessential function of Malpighian tubules is to generate a primary urine via transepithelial fluid secretion, which is mediated by ion and water transport mechanisms in the epithelial cells (principal and stellate cells) of the distal or main segments. The primary urine flows into the proximal or lower segments, which modify the urine (e.g., by ion and/or water reabsorption) before it is emptied into the gut near the junction of the midgut and hindgut. The urine then flows through the ileum and rectum where it can be further modified before it is excreted.

In the context of ion balance, Malpighian tubules are considered the primary tissues for regulating hemolymph calcium (Ca^2+^) concentrations. The tubule epithelium absorbs Ca^2+^ from the hemolymph and either: 1) stores it as mineralized concretions within the cells and/or tubule lumen; or 2) secretes it in soluble form into the lumen for eventual urinary excretion. In *Rhodnius prolixus* (5^th^ stage) and *Acheta domesticus* (adult), nearly all Ca^2+^ that enters the tubules is stored within intracellular concretions; i.e., transepithelial secretion of Ca^2+^ is nominal (Maddrell et al., 1991; Browne and O’Donnell, 2018a). In adult *Drosophila melanogaster*, approximately 85% of the Ca^2+^ that enters the Malpighian tubules is stored as concretions and 15% is secreted, whereas in tubules of larval *D. melanogaster* Ca^2+^ handling is further skewed toward storage (Dube et al., 2000b; Browne and O’Donnell, 2016). The propensity of the tubules to store the majority of absorbed Ca^2+^ is likely widespread among insects given that mineralized Ca^2+^-rich concretions have been observed in lumens and/or epithelial cells of Malpighian tubules from diverse species (Waterhouse, 1950; Clark, 1958; Bradley et al., 1990; Sohal et al., 1976; Maddrell et al., 1991; Wessing et al., 1992; Wessing and Zierold, 1999; Ballan-Dufrançais, 2002; Farina et al., 2022) and a large proportion of the total insect Ca^2+^ content (e.g., 30% in adult *D. melanogaster*, 88% in larval *Drosophila hydei*) often resides in the Malpighian tubules (Wessing and Zierold, 1992; Dube et al., 2000a).

The combined activities of Ca^2+^ storage and secretion provide Malpighian tubules with a remarkable efficiency and capacity to process hemolymph Ca^2+^, especially among Diptera (Browne and O’Donnell, 2018b). Notably, in *D. melanogaster* and *Aedes aegypti*, it is estimated that the tubules can remove the entire hemolymph Ca^2+^ content within as little as 10 min and 18 min, respectively (O’Donnell and Maddrell, 1995; Browne and O’Donnell, 2018b). Accordingly, experiments with flies suggest that Malpighian tubules play key roles in the processing of excess dietary Ca^2+^ that is absorbed into the hemolymph by the midgut. For example, after larval fruit flies (*D. hydei*) were treated with elevated dietary Ca^2+^, the number of type-I concretions, which are mainly composed of mineralized Ca^2+^, increased in the tubule lumen (Wessing and Zierold, 1992). Intriguingly, when adult fruit flies (*D. melanogaster*) were fed high-Ca^2+^ diets, neither their tubular Ca^2+^ content nor Ca^2+^ secretion rates were affected compared to those on standard and low Ca^2+^ diets (Dube et al., 2000a). Moreover, hemolymph Ca^2+^ concentrations were unaffected, suggesting the resting rates of tubular Ca^2+^ secretion were sufficient for excreting the excess Ca^2+^ (Dube et al., 2000a). In another example, when adult blow flies (*Calliphora vicina*) were fed diets with elevated Ca^2+^, the Malpighian tubules showed relative increases of both Ca^2+^ content and uptake rates that were nearly 2-times greater than those found in the entire body (Taylor, 1985).

Previous studies in Diptera suggest that Ca^2+^ stored in Malpighian tubules is not fixed; i.e., it can be mobilized to other tissues depending on the physiological demands of the insect. For example, in female *C. vicina*, decreases of Malpighian tubule Ca^2+^ content were correlated with commensurate increases of ovarian Ca^2+^ content, suggesting tubular Ca^2+^ stores are reabsorbed into the hemolymph for use by the ovaries during reproduction (Taylor, 1984). Along similar lines, in the face fly *Musca autumnalis*, development from larval to pupal stages was associated with progressive decreases in tubular Ca^2+^ that coincided with quantitatively similar increases in cuticular Ca^2+^ content (Krueger et al., 1988). Likewise, in *D. melanogaster* larvae, tubule Ca^2+^ stores are dynamically regulated and proposed to be reabsorbed into the hemolymph for maintaining hemolymph Ca^2+^ levels during periods of rapid growth and pupation (Browne and O’Donnell, 2016). Thus, in the context of Ca^2+^ homeostasis, insect Malpighian tubules appear to perform analogous roles as vertebrate bone.

To date, the cellular and molecular mechanisms of Ca^2+^ transport in insect Malpighian tubules have not been fully elucidated. Electrophysiological evidence from a diversity of insect species indicates that tubular Ca^2+^ uptake from the hemolymph primarily occurs across basolateral membranes of the majority principal cells and not the secondary/stellate cells (Browne and O’Donnell, 2018b). Consistent with this notion, mineralized intracellular concretions have only been observed in principal cells (e.g., Wigglesworth and Salpeter, 1962; Sohal et al., 1976; Ryerse, 1979; Bradley et al., 1990; Wessing and Zierold, 1999; Beyenbach and Piermarini, 2009). A variety of basolateral calcium channels (e.g., transient receptor potential, L-type, cyclic-nucleotide-gated) are hypothesized to be the primary mechanisms of Ca^2+^ uptake from the hemolymph (Dube et al., 2000b; MacPherson et al., 2001; Yu and Beyenbach, 2002; Browne and O’Donnell, 2018a). Cytosolic Ca^2+^ is hypothesized to be stored or sequestered in intracellular compartments by sarco-/endoplasmic reticulum Ca^2+^-ATPases (SERCA) and/or secretory pathway Ca^2+^-ATPases (SPCA) (Rosay et al., 1997; Dube et al., 2000b; Yu and Beyenbach, 2002; Southall et al., 2006; Davies and Terhzaz, 2009; Browne and O’Donnell, 2018a). In addition, cytosolic Ca^2+^ is hypothesized to be extruded into the lumen (secretion) and/or the hemolymph (reabsorption), via apical and/or basolateral plasma membrane Ca^2+^-ATPases (PMCA), respectively (Pannabecker et al., 1995; Dube et al., 2000b). A Na^+^/Ca^2+^ exchange mechanism (e.g., K^+^-dependent Na^+^/Ca^2+^ exchanger, NCKX) has also been hypothesized to contribute to the extrusion of cytosolic Ca^2+^ across the apical or basolateral membranes (Dube et al., 2000b; Overend et al., 2015; Esquivel et al., 2016; Li et al., 2017; Browne and O’Donnell, 2018a).

The goals of the present study are to generate insights into: 1) the role of the Malpighian tubules in the regulation of Ca^2+^ homeostasis in adult female mosquitoes; and 2) the molecular mechanisms of Ca^2+^ transport in Malpighian tubules. To date, few studies have examined the physiological contributions of Malpighian tubules to Ca^2+^ homeostasis in mosquitoes. Mineralized concretions have been detected in principal cells of the distal segments in larval and adult female mosquitoes (Bradley and Nayar, 1985; Bradley and Nayar, 1987; Bradley et al., 1990; Beyenbach and Piermarini, 2009), suggesting a potential role in Ca^2+^ storage. Moreover, high rates of Ca^2+^ uptake have been measured across the basolateral membranes of principal cells in the distal segment of larval *Ae. aegypti* Malpighian tubules (Browne and O’Donnell, 2018b). However, no previous studies have directly examined how Malpighian tubules may contribute to Ca^2+^ homeostasis in adult female mosquitoes.

Using the yellow fever mosquito (*Ae. aegypti*), the present study examined Ca^2+^ regulation in Malpighian tubules of adult females in two independent physiological contexts. First, we measured the Ca^2+^ contents of the Malpighian tubules (and whole mosquitoes) in response to increased Ca^2+^ concentrations of their sucrose diet; in parallel, we quantified the molecular expression of representative putative Ca^2+^ storage (SERCA, SPCA) and Ca^2+^ extrusion (PMCA, NCKX) transport mechanisms in Malpighian tubules. Second, we measured the Ca^2+^ content of Malpighian tubules (and whole mosquitoes) in response to a blood meal; in parallel, we quantified the molecular expression of SERCA, SPCA, PMCA, and NKCX. We hypothesized that Malpighian tubules would accumulate dietary Ca^2+^ (from sugar or blood meals) and that mRNAs encoding Ca^2+^ transporters involved with Ca^2+^ regulation by Malpighian tubules would be differentially expressed in response to dietary Ca^2+^ loading and/or blood feeding.

## 2. Materials and methods

### 2.1. Mosquito Culture and Colony Maintenance

*Ae. aegypti* (Liverpool strain) were reared as previously described (Calkins et al., 2015). Mosquitoes were maintained in a growth chamber set at 27°C, 80% relative humidity, and a 12 h:12 h light:dark cycle. Mosquito eggs were hatched by placing them in 200 ml of distilled water under vacuum for 1 h. After hatching, mosquito larvae were transferred to a 4.4 L rectangular container filled with approximately 2 L of distilled water (∼500 larvae per container) and provided one tropical fish food tablet daily (TetraMin®, Blacksburg, VA, USA). Mosquito pupae were transferred to a 12×12×12” meshed cage and adult *Ae. aegypti* were fed 10% sucrose *ad libitum* after emergence. All experiments were performed on adult female mosquitoes as described in the following sections. When additional eggs were needed, cages of adult females were fed defibrinated rabbit blood purchased from a commercial supplier (Hemostat Laboratories, Dixon, CA, USA) and provided with a coffee filter soaked in distilled water as a site for oviposition.

### 2.2. Dietary Ca^2+^ loading and Sample Collection for Ca^2+^ Content Measurements

#### 2.2.1. Sucrose Feeding

To determine the effects of dietary Ca^2+^ loading via sucrose meals on the Ca^2+^ content of tissues (Malpighian tubules, midguts) or whole bodies of adult females, various concentrations of CaCl_2_ were added to 10% sucrose. In brief, pupae were divided into several 250 ml plastic cups (∼100 pupae/ cup), and each group was transferred into a small cage (32-ounce Ziploc Cup with a screw-on mesh lid). Before eclosion of adults, the cages were provided cotton wicks soaked in 10% sucrose (control) or 10% sucrose containing CaCl_2_ (1, 5, 10, 20, 40, 50, or 80 mM). After 6 days, the sucrose was removed from the cage to allow any recently ingested food to be processed by the mosquitoes (e.g., emptied from the crop and digested/absorbed by the midgut). 24 h after removing the sucrose (on day 7), cages were placed in a refrigerator (4°C) for 5 min to immobilize mosquitoes.

For both the control and CaCl_2_ treated groups, Malpighian tubules and midguts were isolated from 25-30 mosquitoes. Each tissue type from control and CaCl_2_ treated groups was pooled into separate 1.5 ml tubes containing 500 µl of phosphate-buffered saline (PBS; 11.9 mM Phosphates, 137 mM NaCl, and 2.7 mM KCl, pH = 7.4) to represent one biological replicate. The tubes were centrifuged (13,300 *rcf*) at room temperature for 5 min to pellet the tissues; the PBS supernatant was then discarded. In addition, 3-5 intact females were removed from the control and CaCl_2_ treated groups and pooled together in separate 1.5 ml microcentrifuge tubes to represent one biological replicate of the whole body samples. Both the tissue and whole body samples were stored at -20°C until the day of analysis. At least 5 biological replicates (each derived from a unique hatch of mosquito eggs) were obtained for tissue and whole body samples in both the control and CaCl_2_ treated groups.

#### 2.2.2. Blood feeding

One week after emergence, a cage of ∼300 adult female *Ae. aegypti* was starved for 24 h by removing the 10% sucrose solution. Approximately 15 min before providing blood, ∼40 females were removed from the cage to collect samples from a non-blood fed control group (see sample collection details below). The remainder of the mosquitoes were provided access to defibrinated rabbit blood (Hemostat Laboratories, Dixon, CA, USA) via a membrane feeder (Hemotek, Blackburn, UK) as previously described (Calkins and Piermarini, 2017). After a 1 h feeding period, the membrane feeder was removed, and all mosquitoes were immobilized on ice. Mosquitoes with distended abdomens containing visible blood were selected and divided into 6 separate 32-ounce cages containing cotton wicks soaked in 10% sucrose (∼40 females/ cage). Each cage was designated for collection of tissue (Malpighian tubules, ovaries) and whole body samples at a specific time point after the feeding period: 2 h, 6 h, 12 h, 24 h, 48 h, and 72 h. Malpighian tubules and whole body samples were collected and pooled from non-blood fed controls and each time point as described in **2.2.1**. Ovaries were isolated from 20-30 mosquitoes at each time point and pooled together. The pooled tissue and whole body samples were handled and stored as described in **2.2.1**. Five biological replicates (each derived from a unique hatch of mosquito eggs) were obtained for each sample and time point.

### 2.3. Measurement of Ca^2+^ Content

Ca^2+^ contents of tissues and whole bodies were determined by measuring the Ca^2+^ concentrations of homogenates using a Ca^2+^-selective ion meter (HORIBA LAQUAtwin, Kyoto, Japan) calibrated with CaCl_2_ standards (0.13, 0.25, 0.50, 1.25, 2.50, 5.00 mM) dissolved in a Tris-HCl buffer (i.e., 1N HCl neutralized by 2.67 M Tris-Base solution). The standards produced a slope of 28.85 ± 1.0 mV/decade for sample Ca^2+^ concentrations ranging from 0.50 to 5.00 mM (whole bodies and Malpighian tubules; R^2^ = 0.995-1) and 26.33 ± 0.49 mV/decade for concentrations ranging from 0.13 to 1.25 mM (midguts, ovaries, and defibrinated rabbit blood; R^2^ = 0.989-0.997). The tissue or whole body samples were homogenized in 1 N HCl (100 µl per 10 tissues or whole body) manually using plastic pestles (Thermo Fisher Scientific Inc., Waltham, MA, USA). To enhance homogenization and solubilization, the samples were incubated at 70 ℃ for 1 h.

After incubation, an appropriate volume of 2.67 M Tris-Base was added to each tube to reach a final Tris-Base: HCl ratio of 1:2 (v:v); this step neutralizes the pH of the sample solution, which according to the manufacturer (HORIBA) is required for compatibility with the ion meter. The samples were then centrifuged at 13,300 *rcf* for 5 min to pellet insoluble material and 300 µl of the supernatant were applied to the recording chamber of the ion meter. The resulting voltage displacement was recorded. Each sample was measured twice, and the average voltage readings were compared with the standard curve to calculate sample Ca^2+^ concentrations. Ca^2+^ content in nmol was then calculated from the product of the concentration and sample volume.

To measure the Ca^2+^ content of defibrinated rabbit blood, 100 µl of whole blood was homogenized using plastic pestles in 300 µl of 1N HCl and neutralized by 150 µl of 2.67 M Tris-Base. The blood samples were then centrifuged at 13,300 *rcf* for 5 min to pellet insoluble material and 300 µl of the supernatant was applied to the recording chamber of the ion meter. Ca^2+^ concentration and content was then determined as described above.

### 2.4. Capillary Feeder (CAFE) Assay

To estimate the amount of Ca^2+^ ingested by female mosquitoes over six days after feeding on Ca^2+^-loaded sucrose diets, capillary feeder assays were conducted, similar to those described by Corfas and Vosshall (2015). In brief, three newly emerged adult female *Ae. aegypti* were placed in a 28.5 x 95 mm polystyrene *Drosophila* vial (VWR International, Radnor, PA, USA) covered with a cotton plug. Each vial represented one biological replicate. Two 5 µl calibrated capillary tubes (VWR International) filled with 10% sucrose containing CaCl_2_ of various concentrations (see **2.2.1**) were inserted through the cotton plug. Vials with filled capillaries but no mosquitoes were used to record evaporative losses. Every 24 h, the capillaries were removed from vials and replaced with fresh capillaries. The distance (mm) between the starting and ending levels of the removed capillary tubes were measured using a digital caliper (Traceable Carbon Fiber Caliper-4″, VWR International) to determine the volume consumed (after adjusting for evaporative losses). The estimated amount (nmol) of Ca^2+^ ingested per mosquito over 6 days was determined from the product of the total ingested volume and Ca^2+^ concentration of the 10% sucrose.

### 2.5. Sample Collection for Quantitative Real-Time PCR

#### 2.5.1. Sucrose Feeding

Approximately 900 mosquito pupae were divided into three groups (∼300 pupae/group) held in 250 ml plastic cups. Each cup was placed into an 8×8×8” cage. Within 24 h of all adults emerging, cotton wicks containing 10% sucrose (control) or 10% sucrose with CaCl_2_ (5 or 50 mM) were added to the cages. At 24 h (1 day), 72 h (3 days), and 168 h (7 days) after adding the wicks, Malpighian tubules were isolated from ∼30 females of each cage. The Malpighian tubules were pooled according to the treatment into separate 1.5 ml tubes containing PBS and centrifuged (13,300 *rcf*) at 4 ℃ for 5 min. The supernatant was removed and 200 µl of TRIzol™ Reagent (Invitrogen™, Waltham, MA, USA) was added to each tube. The tubes were stored at -80 ℃ until the day of use. A biological replicate consisted of pooled Malpighian tubules from ∼30 females for each dietary Ca^2+^ concentration on each day; a total of 5 biological replicates (derived from 5 independent hatches) were obtained.

#### 2.5.2. Blood feeding

Adult female *Ae. aegypti* were provided blood meals and Malpighian tubules were isolated from ∼30 females as described in **2.2.2**. The Malpighian tubules were pooled, handled, and stored as described in **2.5.1**. A biological replicate consisted of Malpighian tubules from ∼30 females for each time point; a total of 5 biological replicates (derived from 5 independent hatches) were obtained.

### 2.6. Quantification of mRNA expression

Reverse-transcriptase quantitative PCR (qPCR) was used to measure mRNA expression of genes encoding four Ca^2+^-transporters of interest in Malpighian tubules: *SERCA*, *SPCA*, *PMCA*, and *NCKX1*. Total RNA was extracted from isolated Malpighian tubules as previously described (Chomczynski and Sacchi, 1987). In brief, pooled Malpighian tubules were thawed on ice and homogenized in 200 µl TRIzol™ Reagent using sterilized pestles. After homogenization, the total RNA of each sample was extracted using the phenol-chloroform method and then treated with DNase using the Turbo DNA-free™ kit (Invitrogen™). RNA concentration was determined with a spectrophotometer (Nanodrop 2000, Thermo Fisher Scientific Inc.). 200 ng of total RNA from each sample was reverse transcribed into cDNA using random hexamer primers and GoScript™ Reverse Transcriptase kit (Promega Corporation, Madison, WI, USA). The synthesized cDNA was stored at -20 ℃.

Primers to amplify approximate 100 bp regions of the Ca^2+^-transporter cDNAs were designed using Primer3 software (Table S1; Untergasser et al., 2012). Primers that amplify two reference genes (*actin* and *RPS17*) established by Dzaki et al. (2017) were also utilized. All primers were synthesized by Integrated DNA Technologies (IDT Inc., Morrisville, NC, USA). Standard curves were used to measure the efficiency of primer amplification for each gene of interest, which ranged from 98% to 102%. qPCR was performed in 96-well Multiplate™ PCR plates (Bio-Rad Inc., Hercules, CA, USA) on a Real-time PCR thermocycler (Bio-Rad CFX96 Touch) at The Ohio State University (OSU) Molecular and Cellular Imaging Center (MCIC) on the OSU Wooster campus (Wooster, OH). Each 10 µl PCR reaction consisted of 5 µl of GoTaq® qPCR Master Mix (Promega Corporation), 0.5 µl of 10 µM forward primer, 0.5 µl of 10 µM reverse primers, and 2 µl of cDNA template. Each PCR cycle started with an initial denaturation at 95 ℃ for 2 min, followed by 39 cycles of 95°C (20 s), 55°C (20 s), and 68°C (20 s). A melt curve was run at the end of each cycle to ensure that only a single product was amplified in each well of the qPCR plate. The relative mRNA expression results were calculated using the 2^-ΔCt^ method and normalized to the average Ct value of the two reference genes.

### 2.7. Statistical analyses

R (version 4.1.2), RStudio (version 2022.02.3, RStudio Inc., Boston, MA, USA), or Prism (version 9; GraphPad Software Inc., La Jolla, CA, USA) were used for all statistical analyses. Shapiro normality tests were conducted for each dataset; any non-normal datasets were log transformed before analysis. Data figures were generated with Prism.

#### 2.7.1. Measurement of Ca^2+^ content

For sucrose Ca^2+^ loading trials, the mean Ca^2+^ content of Malpighian tubules or whole bodies from mosquitoes fed Ca^2+^-treated sucrose were compared to controls using a one-way analysis of variance (ANOVA) with a Holm-Sidak post hoc test (multiple comparisons were only made versus the control; p < 0.05). For blood feeding trials, the mean Ca^2+^ content of Malpighian tubules or whole bodies at each time point before and after a blood meal was compared using a randomized block one-way ANOVA and Fisher’s least significant difference (LSD) post hoc tests (p < 0.05).

#### 2.7.2. Quantification of mRNA expression

For dietary-Ca^2+^ loading trials, the mean mRNA expression levels of each gene of interest on each day were compared between control and treatment groups using randomized block one-way ANOVA and Fisher’s LSD post hoc tests. For blood feeding trials, the mean mRNA expression levels of each gene of interest were compared between each time point before and after blood feeding using a randomized block one-way ANOVA and Fisher’s LSD post hoc tests (p < 0.05).

## 3. Results

### 3.1. Sucrose Ca^2+^ loading

#### 3.1.1. Ca^2+^ content

In adult female *Ae. aegypti* fed on 10% sucrose with nominal Ca^2+^ (controls) for 6 days, the Ca^2+^ content of the Malpighian tubules (i.e., in all 5 tubules) per mosquito was 31.2 ± 2.1 nmol (n = 16), representing ∼18% of the whole body Ca^2+^ (172.7 ± 8.5 nmol/mosquito; n = 15). As shown in Figure 1A, when mosquitoes were fed 10% sucrose containing elevated concentrations of CaCl_2_ (referred to as Ca^2+^ treatments hereafter) for 6 days, the Ca^2+^ contents of the Malpighian tubules were elevated (P < 0.05) compared to the controls in all treatments except for 1 mM Ca^2+^. The increase of Ca^2+^ content appeared to be concentration-dependent and plateau by the 10 mM Ca^2+^ treatment (Figure 1A). Similarly, whole body Ca^2+^ contents were greater (P < 0.05) in the Ca^2+^ treatments compared to the controls, except for the 1 mM Ca^2+^ treatment (Figure 1B). The increase of whole body Ca^2+^ appeared to be less concentration-dependent than that in the Malpighian tubules and plateau by the 5 mM Ca^2+^ treatment (Figure 1B). The percentage of whole body Ca^2+^ attributed to the Malpighian tubules increased (P < 0.05) in the Ca^2+^ treated groups (except for 1 mM Ca^2+^) relative to controls, reaching as much as ∼38% (Figure S1). Among the Ca^2+^ treatments causing significant effects (i.e., 5-80 mM), the Malpighian tubules were responsible for ∼37-70% of the whole body increase in Ca^2+^ (Table S2).

**Figure 1.**
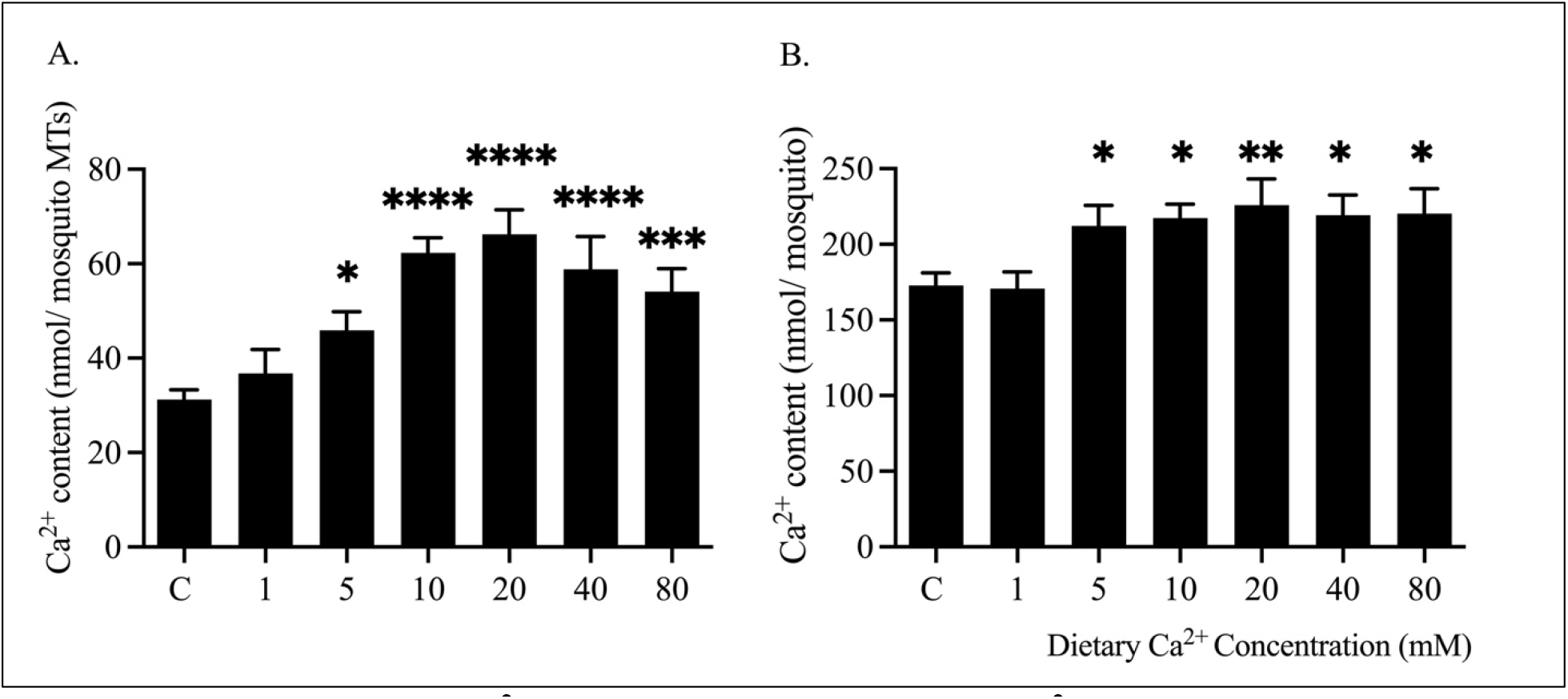
Effects of dietary Ca^2+^ treatments on the content of Ca^2+^ in Malpighian tubules (MTs) (A) and whole bodies (B) of adult females. Asterisks indicate significant differences between control (C) and each treatment group as determined by ordinary one-way ANOVA and Holm-Sidak post hoc tests (p < 0.05). Values are mean ± s.e.m. For Malpighian tubules, n = 16, 5, 7, 7, 8, 8, and 7, respectively, for C, 1, 5, 10, 20, 40, and 80 mM Ca^2+^ treatments; for whole bodies, n = 15, 5, 7, 7, 8, 7, and 6, respectively, for C, 1, 5, 10, 20, 40, and 80 mM Ca^2+^ treatments.

To provide a point of comparison for the Malpighian tubules, we also attempted to measure the Ca^2+^ contents of midguts isolated from mosquitoes in the control and Ca^2+^ treatment groups. In both mosquitoes from the control and majority of Ca^2+^ treatments, midgut Ca^2+^ contents were below detectable limits of the Ca^2+^ ion meter (i.e., < 1.49 nmol per midgut). However, Ca^2+^ contents of midguts from mosquitoes in the 40 and 80 mM Ca^2+^ treated mosquitoes were measurable, which were 3.41 ± 0.54 (n = 5) and 6.73 ± 0.99 nmol of Ca^2+^ per midgut (n = 5) respectively.

To provide better physiological context for the increases in whole body Ca^2+^ content measured during the Ca^2+^ treatments (5-80 mM), we performed separate experiments to estimate the amount of Ca^2+^ ingested in each Ca^2+^ treatment. As shown in Table S3, the estimated amount of ingested Ca^2+^ exceeds the increases in the whole bodies, especially during the 10-80 mM Ca^2+^ treatments. As explained in the Discussion **4.1**, we interpret the differences between these values as an estimate of the amount of Ca^2+^ excreted by the mosquitoes over the experimental period (Table S3).

#### 3.1.2. mRNA expression

To provide insights into putative molecular mechanisms contributing to the changes in Malpighian tubule Ca^2+^ content associated with dietary Ca^2+^ concentration (Figure 1A), we measured mRNA expression levels of select Ca^2+^-transporters hypothesized to contribute to intracellular sequestration (SERCA, SPCA) or extrusion (PMCA, NCKX1) of cytosolic Ca^2+^. In these experiments we fed mosquitoes 10% sucrose with 5 mM or 50 mM CaCl_2_, which correspond to treatments expected to result in intermediate or maximal effects on tubule Ca^2+^ content, respectively (Figure 1A); control mosquitoes were fed 10% sucrose without CaCl_2_. The mRNA expression levels of each gene in tubules of controls were stable among days 1, 3, and 7 of the experiments (Figure S2). Quantiatively, *NCKX1* was the most abundant mRNA, followed by *PMCA, SERCA*, and *SPCA* (Figure S2).

As shown in Figure 2, *SERCA* mRNA levels in Malpighian tubules of both the 5 mM and 50 mM Ca^2+^ treatments were up-regulated relative to those in the controls on day 3, but the change was relatively small (∼1.2-fold higher; Figure 2A). *SPCA* mRNAs in Malpighian tubules of the 50 mM Ca^2+^ treatment were elevated slighty compared to those of controls on days 1, 3, and 7 by ∼1.4-1.5 fold, whereas in those of 5 mM Ca^2+^ treatment, the mRNA levels of *SPCA* remained unchanged relative to controls on day 1, but was up-regulated to a similar level as those of 50 mM Ca^2+^ treatment on days 3 and 7 (Figure 2B). At each time point, *PMCA* mRNA levels in Malpighian tubules were similar between the controls and 5 mM Ca^2+^ treatment (Figure 2C). However, the mRNA abundances of *PMCA* in Malpighian tubules of the 50 mM Ca^2+^ treatment were ∼2-fold higher than those of the controls and the 5 mM Ca^2+^ treatments on days 1, 3, and 7 (Figure 2C). The mRNA levels of *NCKX1* in Malpighian tubules were similar to controls among the treatments at each time point, except for day 1 when the *NCKX1* mRNA abundance was ∼1.5 fold up-regulated in tubules of 50 mM Ca^2+^ treatment compared to those of control and 5 mM Ca^2+^ treated groups (Figure 2D).

**Figure 2.**
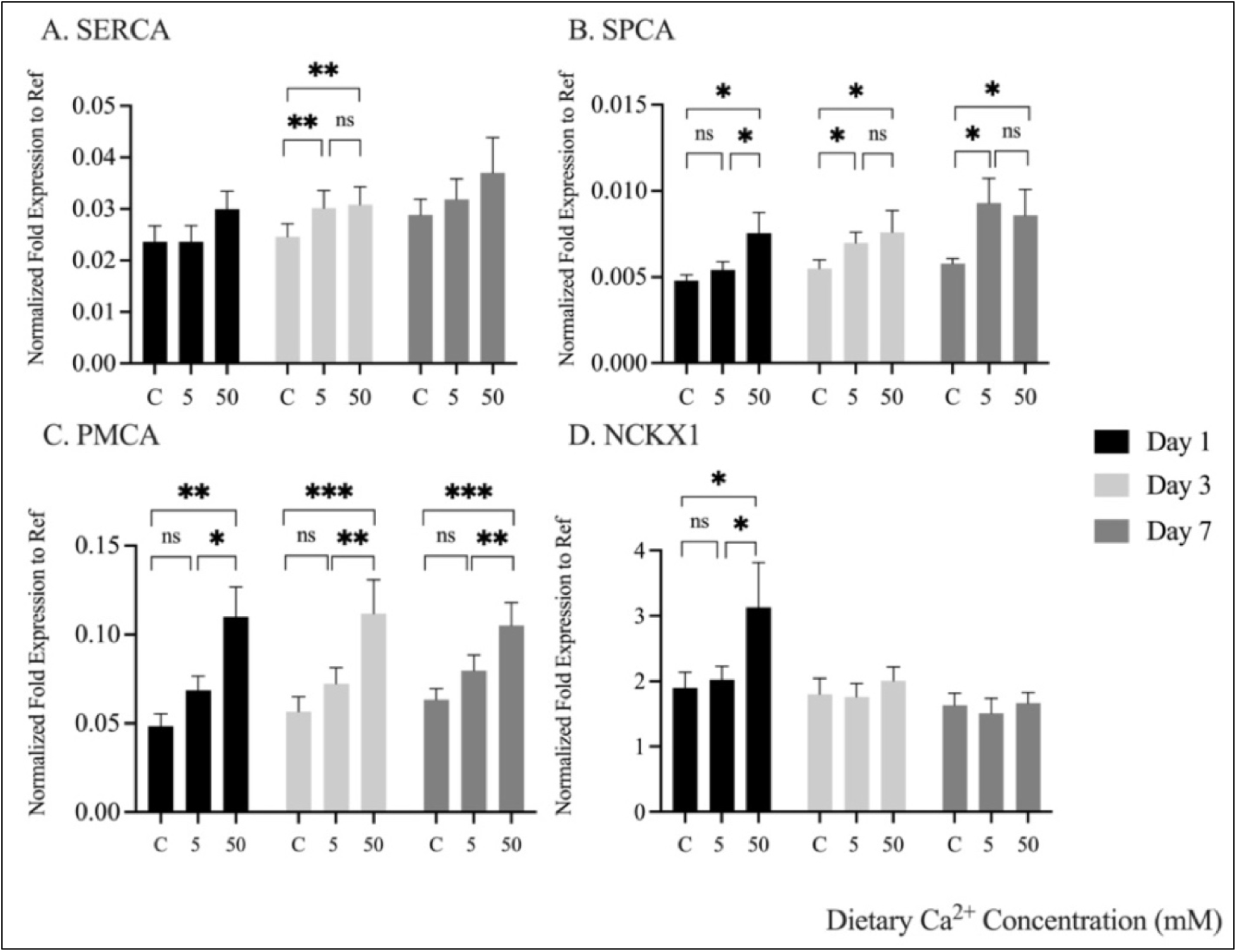
Effects of dietary Ca^2+^ treatments on *SERCA* (A)*, SPCA* (B)*, PMCA* (C), and *NCKX1* (D) mRNA levels in *Ae. aegypti* Malpighian tubules at days 1, 3, and 7 (indicated by black, light grey, and dark grey bars separately). Asterisks indicate differences among means within a time point as determined by a randomized block one-way ANOVA and Fisher’s LSD tests (p < 0.05). C = control. Values are mean ± s.e.m; n = 5 replicates.

### 3.2. Blood Feeding

#### 3.2.1. Ca^2+^ content

In non-blood fed adult female *Ae. aegypti* starved for 24 h (control), the Ca^2+^ content of the Malpighian tubules per mosquito (i.e., in all 5 tubules) was 29.51± 0.81 nmol (n = 5), representing ∼22% of the whole body Ca^2+^ content. In blood fed *Ae. aegypti*, the Ca^2+^ contents of Malpighian tubules changed dynamically over the next 72 h (Figure 3A). The Ca^2+^ content first slightly increased by ∼4 nmol to 33.38 ± 0.95 nmol/mosquito tubules by 6 h post-blood feeding (PBF). To put this increase in context, the estimated amount of ingested Ca^2+^ from a blood meal was ∼3.6 nmol, assuming a typical consumption of ∼3 µl of blood/female (Zhou et al., 2007) and the measured Ca^2+^ concentration of defibrinated rabbit blood (∼1.2 mM; see **2.2.3**).

**Figure 3.**
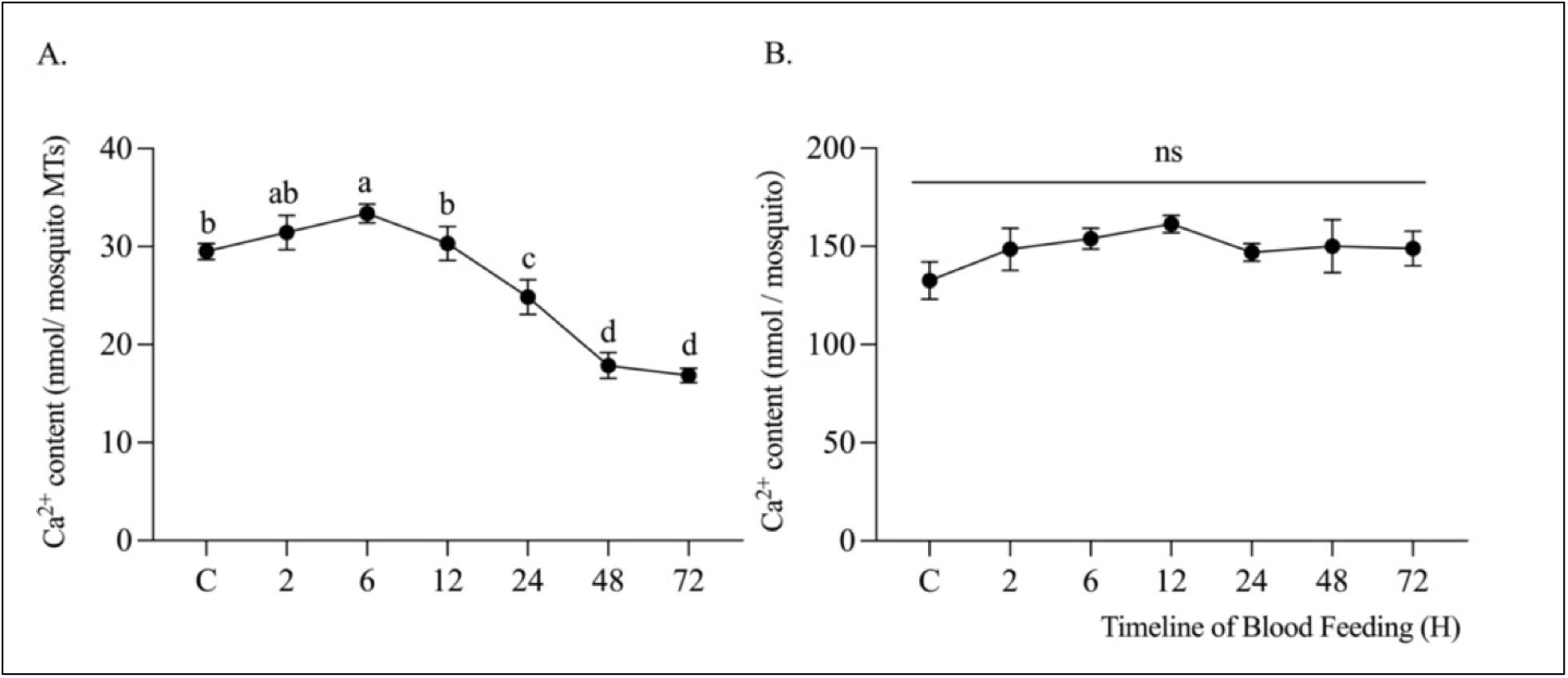
Effects of blood feeding on the content of Ca^2+^ in Malpighian tubules, MTs (A) and whole bodies (B). Lower-case letters indicate statistical categorizations of the means as determined by a randomized block one-way ANOVA and Fisher’s LSD tests (p < 0.05). Values are mean ± s.e.m; n = 5 replicates.

Between 6-48 h PBF, the tubule Ca^2+^ content steadily decreased, reaching 17.85 ± 1.32 nmol/mosquito tubules by 48 h PBF (Figure 3A). This value is nearly 50% lower than tubules from control mosquitoes. At 72 h PBF, the Ca^2+^ content of the tubules was similar to that at 48 h PBF (Figure 3A). In contrast, the Ca^2+^ contents of whole bodies of blood-fed mosquitoes were similar to those of non-blood fed controls at each time point sampled (Figure 3B). Thus, from 48-72 h PBF the tubules only represented ∼11% of the whole body Ca^2+^ content. The prominent decrease of tubule Ca^2+^ content at 48-72 h PBF raised the question of whether it was being redistributed to the ovaries as found in *C. vicina* during oogenesis (Taylor, 1984). Ca^2+^ contents of the ovaries were below detectable limits (i.e., < 1.47 nmol per mosquito ovaries) in the control and majority of PBF time points, but at 48 h and 72 h PBF the Ca^2+^ content of the ovaries was measurable (2.82 ± 0.29 nmol per mosquito ovaries; n = 5).

#### 3.2.2. mRNA expression

To provide insights into putative molecular mechanisms contributing to the dynamic changes in Ca^2+^ content of Malpighian tubules after blood feeding (Figure 3), we measured the transcript abundance of *SERCA, SPCA, PMCA*, and *NCKX1*. The mRNA abundance of *SERCA* showed a progressive up-regulation from 2-6 h PBF reaching levels ∼3-fold higher than controls (Figure 4A). This elevated expression level of *SERCA* persisted until 24 h PBF and returned to those of controls by 72 h PBF (Figure 4A). Similary, *SPCA* mRNA levels were up-regulated (∼1.5-fold) between 6-24 h PBF and then returned progressively to control levels by 72 h PBF (Figure 4B). Within 2 h post-blood feeding (PBF), the mRNA abundance of *PMCA* in Malpighian tubules was up-regulated (∼2-fold) compared to controls and remained up-regulated through 48 h PBF (Figure 4C). Between 48-72 h PBF, *PMCA* mRNA levels returned to those of controls. In contrast, differential expression of *NCKX1* mRNA levels was only detected at 24 h PBF when it was ∼1.5-fold higher than MTs of control mosquitoes (Figure 4D).

**Figure 4.**
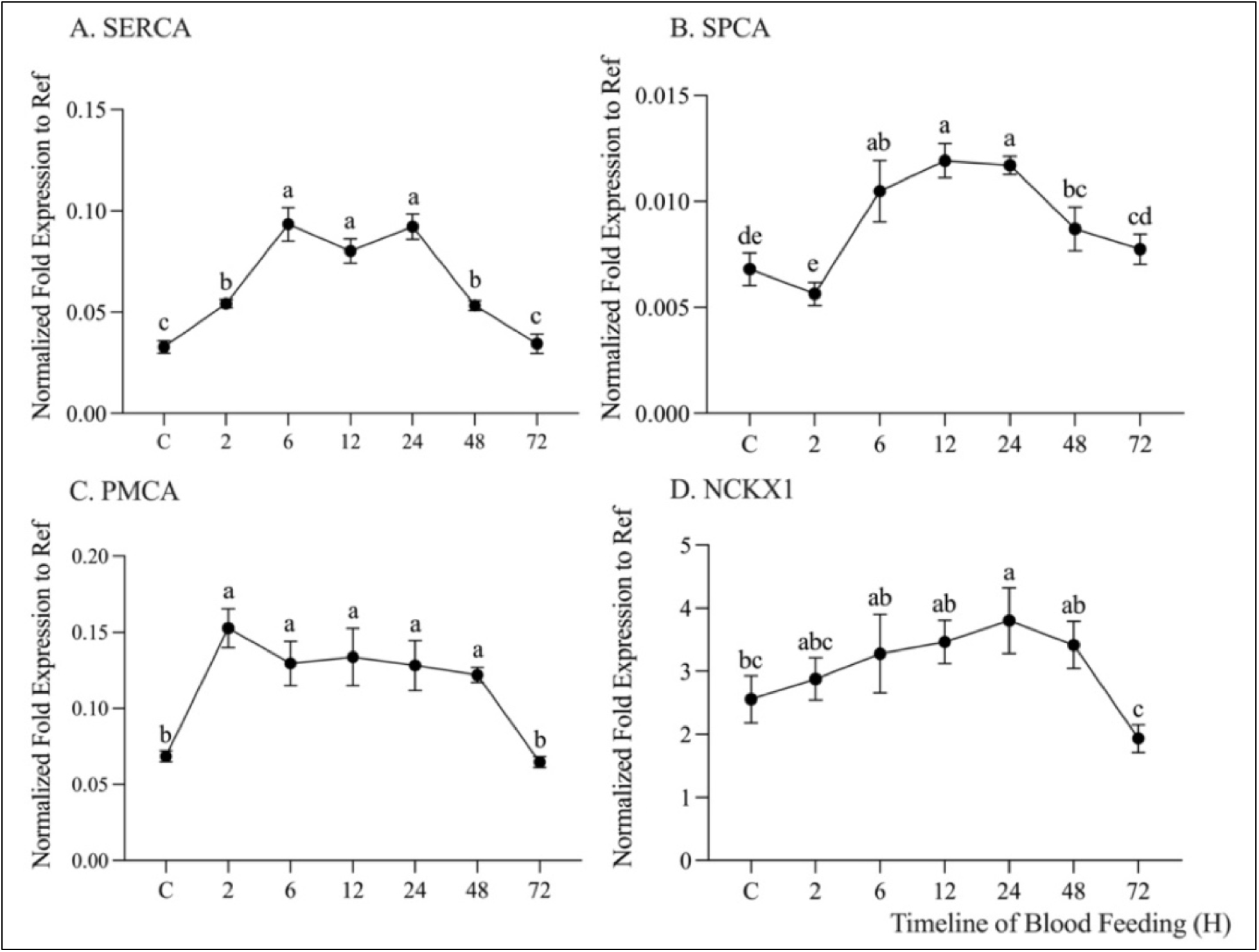
Effects of a blood meal on *SERCA* (A)*, SPCA* (B)*, PMCA* (C), and *NCKX1* (D) mRNA abundance in *Ae. aegypti* Malpighian tubules before (C) and at several time points post blood feeding. The relative mRNA expressions of *SERCA*, *SPCA*, and *PMCA* were log-transformed to fit the assumption of normality before statistical analysis (for clarity, data shown are not log transformed). Lower-case letters indicate statistical categorizations of the means as determined by a randomized block one-way ANOVA and Fisher’s LSD tests (p < 0.05). Values are mean ± s.e.m; n = 5 replicates.

## 4. Discussion

The present study is the first in adult female mosquitoes to elucidate how dietary Ca^2+^ loads and blood feeding affect the Ca^2+^ content of Malpighian tubules and the molecular expression of Ca^2+^ transport mechanisms in mosquito Malpighian tubules. Collectively, the results advance our basic knowledge of the physiological roles that Malpighian tubules play in mosquito Ca^2+^ homeostasis and putative transport mechanisms involved with the renal handling of Ca^2+^ in insects. As discussed in **4.2** below, our study also reveals potential novel roles of the tubules in Ca^2+^ redistribution in mosquitoes during blood meal processing and egg development.

### 4.1. Dietary Ca^2+^ loading via sucrose

When fed 10% sucrose with nominal Ca^2+^, Malpighian tubules of adult female *Ae. aegypti* contain ∼18% of the whole body Ca^2+^ content. Presumably, most of the tubule and whole body Ca^2+^ in adult females is derived from Ca^2+^ that was ingested during the larval stages (Barkai and Williams, 1983). Supplementing the 10% sucrose with CaCl_2_ resulted in increases of the Ca^2+^content of both Malpighian tubules and whole bodies, with the tubules responsible for the majority of the whole body increases in most of the treatments. Based on previous studies in blow flies (*C. vicina*), it is assumed that the majority of Ca^2+^ ingested by insects is absorbed into the hemolymph via the midgut and then absorbed by the tubules for storage and/or excretion (Taylor, 1986). When mosquitoes were fed sucrose containing ≥ 5 mM Ca^2+^, the increases of whole body Ca^2+^ levels were slightly (5 mM Ca^2+^ treatment) or dramatically (10-80 mM Ca^2+^ treatments) below the estimated amounts of Ca^2+^ ingested. We hypothesize that the remaining ingested Ca^2+^ that is unaccounted for in the whole bodies was excreted by the Malpighian tubules (Table S3). Taken together, these findings suggest that adult female mosquitoes retain a small portion of the excess dietary Ca^2+^ in the Malpighian tubules and other tissues, but once the carrying capacity of these tissues is exceeded then Ca^2+^ is excreted. In other dipterans, Malpighian tubules have shown similar responses to enhanced dietary Ca^2+^; i.e., the excess is stored and/or excreted by Malpighian tubules (Taylor, 1985; Wessing and Zierold, 1992; Dube et al., 2000a).

The above results have potential implications for adult female *Ae. aegypti* in natural conditions. That is, when not blood feeding, adult female mosquitoes rely on nectar or other plant juices for nutrition. A survey of nectar samples from 8 flower species determined that the Ca^2+^ concentrations ranged from 0.75-13 mM (McLellan, 1975). Assuming that adult female *Ae. aegypti* ingest flower nectar in similar quantities as 10% sucrose, our results suggest that Malpighian tubules have the capacity to handle dietary Ca^2+^ loads that they would receive from floral sources via storage and/or excretion.

Although Malpighian tubules stored large percentages of the whole body Ca^2+^ content (Figure S1), especially in the elevated dietary Ca^2+^ treatments, other tissues might also contribute to Ca^2+^ storage. For example, in other insects, such as adult stink bugs *Brontocoris tabidus* (Guedes et al., 2007) and ants (Ballan-Dufrançais, 2002), midguts accumulate Ca^2+^-containing granules in columnar cells. Consistent with this possibility, we found that Ca^2+^ contents of midguts became detectable after mosquitoes fed on high Ca^2+^ treatments (40 or 80 mM Ca^2+^). However, compared to the Malpighian tubules, the midgut Ca^2+^ levels at their highest (i.e., during the 80 mM Ca^2+^ treatment) were only ∼12% of that of the Malpighian tubules and represented only ∼3% of the whole body Ca^2+^ content.

The increased Ca^2+^ content of Malpighian tubules observed in the present study likely corresponds to enhanced storage of mineralized Ca^2+^ within principal cells and/or tubule lumens (Maddrell et al., 1991; Wessing and Zierold, 1992; Dube et al., 2000a). Consistent with potential storage in principal cells, we found that Malpighian tubules of mosquitoes fed moderate (5 mM) or high (50 mM) Ca^2+^ diets up-regulated both *SERCA* and *SPCA* mRNA levels. This suggests that in response to elevated dietary Ca^2+^, the tubules increase their molecular potential for transporting cytosolic Ca^2+^ into intracellular storage compartments. The above interpretations are consistent with previous studies in Malpighian tubules that suggest SERCA and SPCA contribute to intracellular Ca^2+^ sequestration (Southall et al., 2006; Browne and O’Donnell, 2018a).

As previously mentioned, when fed high Ca^2+^ diets, mosquitoes likely enhance the renal excretion of Ca^2+^. Consistent with this notion, we found that Malpighian tubules of mosquitoes up-regulated *PMCA* mRNAs when fed high Ca^2+^ diets. Notably, previous studies in lepidopteran Malpighian tubules found PMCA-like immunoreactivity was more abundant in the apical vs. basolateral membranes of principal cells (Pannabecker et al., 1995). Assuming PMCA has a similar localization pattern in principal cells of mosquito Malpighian tubules, an up-regulation of apical PMCA might enhance the molecular capacity of Malpighian tubules for extruding Ca^2+^ across the apical membranes into the lumen for urinary excretion.

*NCKX1* mRNA expression in Malpighian tubules was highly abundant compared to *PMCA*, *SERCA*, and *SPCA* mRNAs, consistent with findings from previous transcriptomic studies (Overend et al., 2015; Esquivel et al., 2016; Li et al., 2017; Hixson et al., 2022). The localization of NCKX1 protein in Malpighian tubules of any insect is unknown, but similar to PMCA, Na/Ca exchange mechanisms have been hypothesized to play roles in extrusion of Ca^2+^ across both the apical and basolateral membranes (Esquivel et al., 2016; Li et al., 2017; Browne and O’Donnell, 2018a). In this study, we found that *NCKX1* mRNA levels in mosquito Malpighian tubules were transiently up-regulated on day 1 in response to the high dietary Ca^2+^ treatment before returning to control levels on days 3 and 7. Thus, it is possible that NCKX1 is specialized for enhancing Ca^2+^ excretion during the acute stages of the high Ca^2+^ treatment, presumably via apical secretion. Figure 5 presents a hypothetical model of Ca^2+^ transport in mosquito Malpighian tubules that illustrates our interpretations of the above molecular changes in response to elevated dietary Ca^2+^ from sucrose meals.

**Figure 5.**
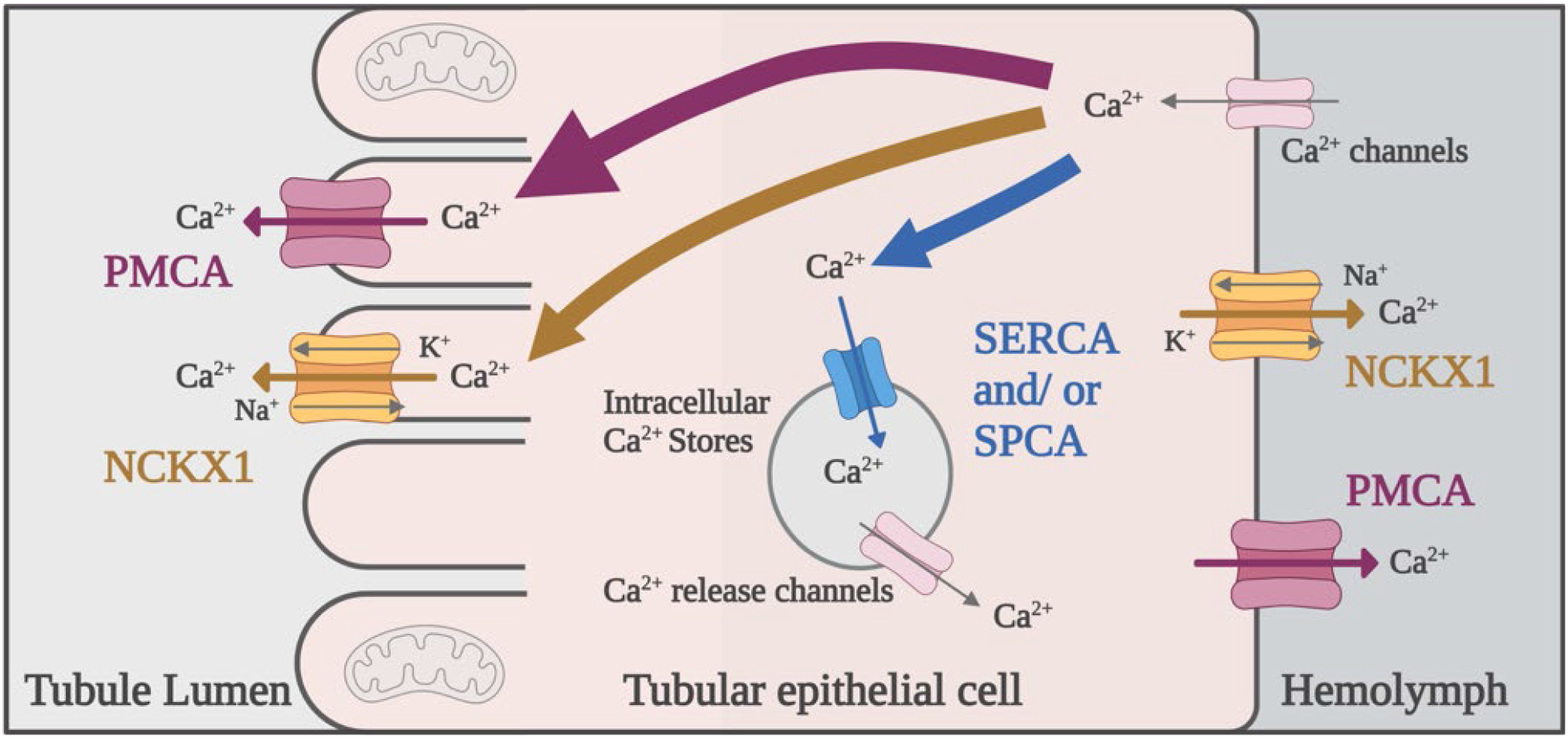
Hypothesized model of Ca^2+^ transporting mechanisms in mosquito Malpighian tubule epithelial cells when fed 10% sucrose with elevated Ca^2+^. Putative localization of transporters is adopted from the Browne and O’Donnell (2018a) model of Ca^2+^ transport in *A. domesticus*. Magenta, yellow, and blue arrows represent the possible Ca^2+^ transport pathways mediated by apical/basolateral PMCA, apical/basolateral NCKX1, and intracellular SERCA/SPCA, respectively. Grey arrows represent putative mechanisms of Ca^2+^ influx (Ca^2+^ channels) and intracellular release (Ca^2+^ release channels) not examined in the present study. Thickened arrows represent our interpretations of the up-regulated mRNAs in Figure 2.

### 4.2. Blood feeding

Our study is the first to provide insights into how the Ca^2+^ content of Malpighian tubules in mosquitoes is affected during blood meal processing and egg development. Notably, the Ca^2+^ content of Malpighian tubules was dynamic within 72 h after ingesting blood and appeared to consist of 3 distinct phases. The first phase (0-6 h PBF) was characterized by a small increase of Ca^2+^ that was remarkably similar to the estimated amount of Ca^2+^ in the ingested blood (i.e., ∼3.6 nmol), suggesting the Malpighian tubules are a sink for nearly all of the ingested Ca^2+^ in the blood meal. Similarly, in the obligate-hematophagous *R. prolixus*, most of the Ca^2+^ from ingested blood was deposited in the upper segments of Malpighian tubules (Maddrell et al., 1991).

The second phase (6-48 h PBF) was characterized by a dramatic decrease in the Ca^2+^ content of the tubules by ∼50%, whereas the third phase (48-72 h PBF) was characterized by a stasis of tubule Ca^2+^ levels. Given that the whole body Ca^2+^ content did not change during phase 2, the decrease in tubule Ca^2+^ content likely reflects its redistribution from mineralized stores to other tissues rather than excretion from the mosquito. One potential site for redistribution of tubule Ca^2+^ is the ovaries (Taylor, 1984). Consistent with this notion, ovarian Ca^2+^ levels were not detectable until the end of phase 2 and remained stable through phase 3, accounting for ∼20% of the decreased tubule Ca^2+^ content. The tissues responsible for the remaining ∼80% of the decreased tubule Ca^2+^ content remain to be determined.

The timing of reduced tubule and increased ovarian Ca^2+^ contents overlaps with vitellogenesis in *Ae. aegypti*, which typically peaks between 24-30 h PBF and ends by 48 h PBF (Dhadialla et al., 1992). Ca^2+^ plays an important role in the signaling of ovaries during vitellogenesis (Dhadialla and Raikhel, 1991; Sappington and S. Raikhel, 1998; Warrier and Subramoniam, 2002). Thus, the changes in tubule and ovarian Ca^2+^ contents during phase 2 may reflect an important role of tubule Ca^2+^ stores in maintaining hemolymph Ca^2+^ levels to accommodate the demands of the ovaries and other tissues that undergo intensive Ca^2+^ cell signaling during vitellogenesis. At least in vertebrate systems, cells undergoing intensive intracellular Ca^2+^ signaling have been shown to decrease the Ca^2+^ concentrations of the surrounding extracellular fluid (Hofer, 2005).

Our qPCR results revealed that the dynamic changes in Ca^2+^ content after blood feeding coincided with dynamic changes in Ca^2+^-transporter mRNA expression. That is, *SERCA* and *SPCA* were up-regulated during phase 1, suggesting the tubules enhance their molecular potential to store intracellular Ca^2+^ as ingested Ca^2+^ is absorbed into the hemolymph. On the other hand, SERCA and SPCA were down-regulated during phases 2 and 3, suggesting the tubules decrease their molecular potential to store intracelluar Ca^2+^ while tubule Ca^2+^ stores are being reduced for use by other tissues (e.g., ovaries).

*PMCA* transcripts showed a similar pattern of up-regulation as *SERCA* and *SPCA*, but remained up-regulated throughout phase 2, before returning to control levels by phase 3. On the other hand, *NCKX1* transcripts showed a gradual up-regulation that did not peak until the middle of phase 2 (24 h PBF), followed by a down-regulation to control levels by the end of phase 3. Taken together, these findings suggest the tubules maximize their molecular potential to extrude cytosolic Ca^2+^ during phase 2, consistent with the 50% decrease in tubule Ca^2+^ content. Given that tubule Ca^2+^ is more likely redstributed to other tissues rather than excreted during phase 2, we hypothesize that the up-regulation of *PMCA* and *NCKX1* mRNAs leads to enhanced basolateral PMCA and NCKX1 protein abundances in Malpighian tubules, which would be expected to enhance the reabsorption of cytosolic Ca^2+^ (presumably released from intracellular stores) into the hemolymph during phase 2 for eventual use by other tissues (e.g., ovaries). Moreover, the down-regulation of putative basolateral PMCA and NCKX1 by phase 3 would coincide with reduced demand/use of tubule Ca^2+^ by other tissues (e.g., ovaries) post-vitelleogensis. Figure 6 presents a hypothetical model of Ca^2+^ transport in mosquito Malpighian tubules that illustrates our interpretations of the above molecular changes after blood feeding during phases 1 and 2.

**Figure 6.**
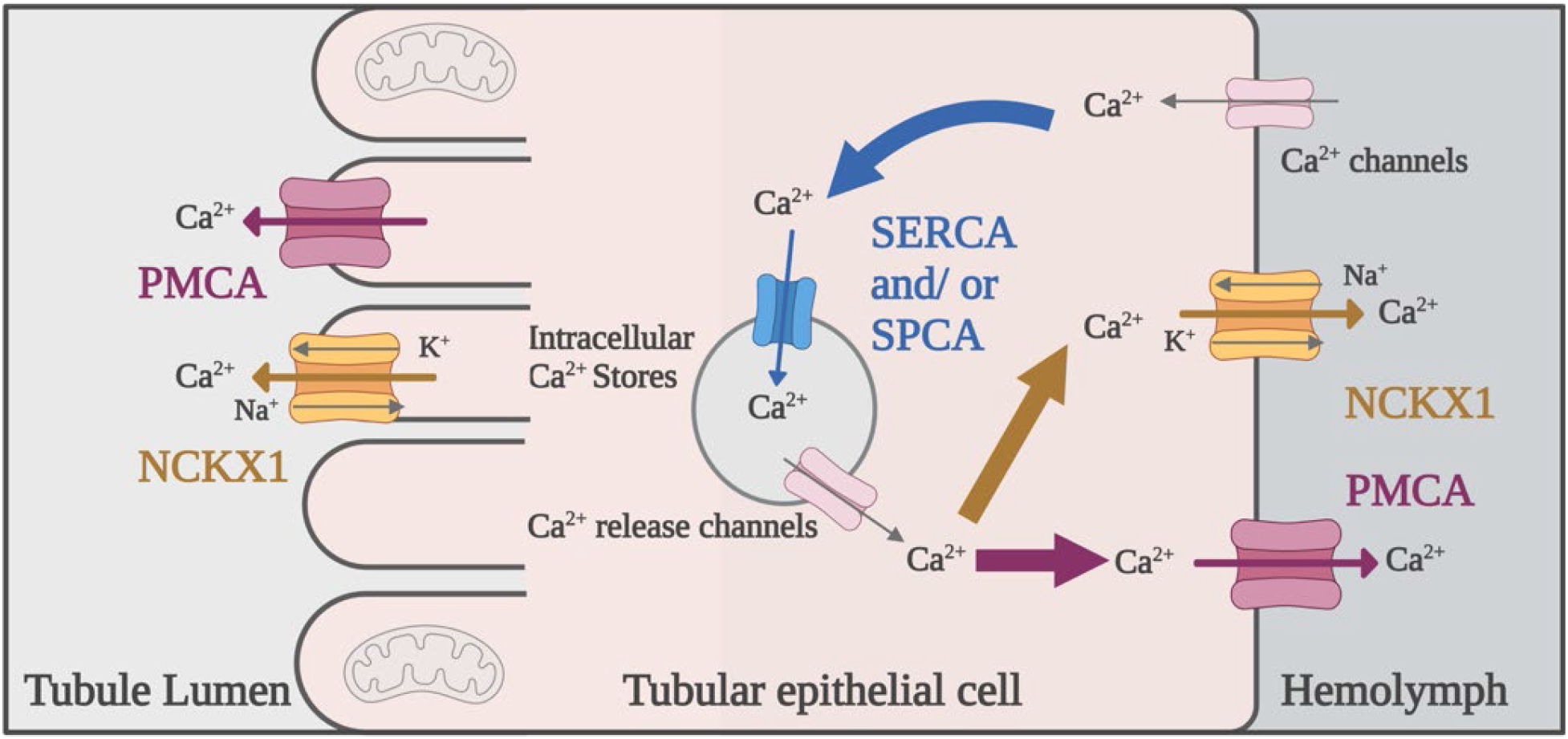
Hypothesized model of Ca^2+^ transporting mechanisms in mosquito Malpighian tubule epithelial cells during Phase 1 and 2 (i.e., 0-48 h) PBF. Putative localization of transporters is adopted from the Browne and O’Donnell (2018a) model of Ca^2+^ transport in *A. domesticus*. Magenta, yellow, and blue arrows represent the possible Ca^2+^ transport pathways mediated by apical/basolateral PMCA, apical/basolateral NCKX1, and intracellular SERCA/SPCA, respectively. Grey arrows represent putative mechanisms of Ca^2+^ influx (Ca^2+^ channels) and intracellular release (Ca^2+^ release channels) not examined in the present study. Thickened arrows represent our interpretations of the up-regulated mRNAs in Figure 4.

In conclusion, the results of our study suggest that Malpighian tubules have a remarkable capacity to handle excess dietary Ca^2+^ in sucrose meals, most likely by first storing Ca^2+^ (via up-regulated intracellular *SERCA* and *SPCA*) and then secreting it (via up-regulated apical *PMCA* and *NCKX1)* once storage capacity is reached. We also found that after ingesting blood the Ca^2+^ levels in Malpighian tubules are dynamic, likely responding to metabolic demands for Ca^2+^ in other tissues (e.g., ovaries) by modulating the molecular capacity of the tubules for intracellular storage (*SERCA*, *SPCA*) and basolateral reabsorption (*PMCA*, *NCKX1*) of Ca^2+^. Future studies will be required to confirm the hypothesized localizations of the above Ca^2+^ transport mechanisms and functional changes in Malpighian tubules in response to elevated dietary Ca^2+^ treatments or blood feeding. Given the indispensable roles of Ca^2+^ in insect physiology, additional studies investigating Ca2+ processing by Malpighian tubules and their underlying Ca^2+^ transport mechanisms may also reveal novel biochemical targets for developing insecticides that disrupt the renal regulation of Ca^2+^ homeostasis in mosquitoes.

## 5. Acknowledgements

The authors thank Dr. Ellen Klinger (OSU) and Dr. Tae Young Lee (OSU) for providing guidance on statistical analyses, Dr. Ana Trabanino (OSU) for assistance with RT-qPCR expreiments, and Dr. Mike O’Donnell (McMaster University) for helpful discussions and insights.

## 6. Appendices

**Table S1.**
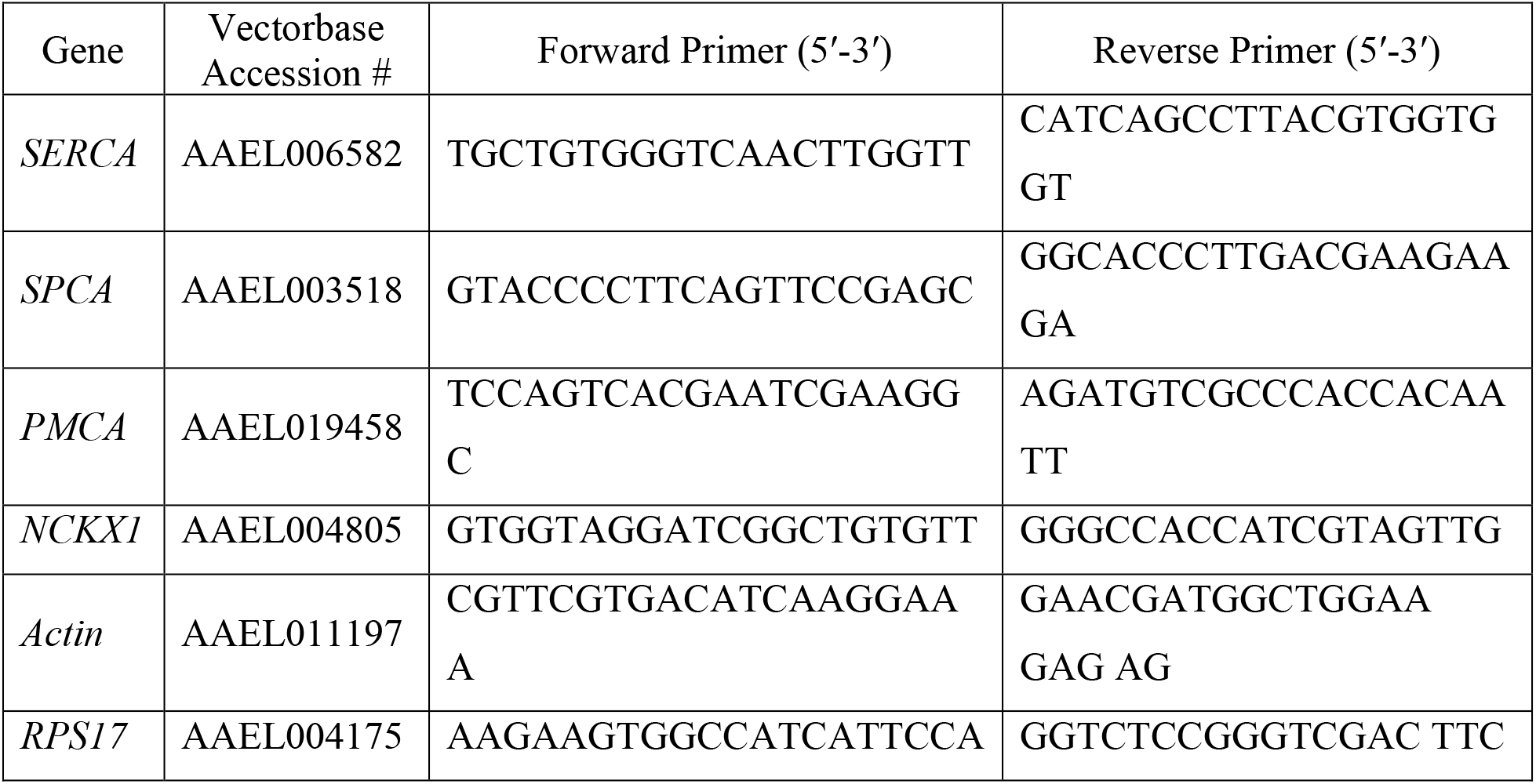
Forward and reverse primers used in qPCR.

**Table S2.** Effects of dietary Ca^2+^ in 10% sucrose on the Ca^2+^ content increases in mosquito whole bodies and Malpighian tubules (data are calculated from Figure 1; values are mean ± s.e.m), and the percentage of the whole body Ca^2+^ increase attributed to Malpighian tubules. Data are only shown for Ca^2+^ treatments that resulted in significant increases to whole bodies and Malpighian tubules (see Figure 1).

| CaCl<sub>2</sub> Concentration<br>(mM) in 10% sucrose | Whole body<br>$\text{Ca}^{2+}$ increase<br>(nmol) | Malpighian tubule<br>$\text{Ca}^{2+}$ increase<br>(nmol) | Percentage of whole body<br>$\text{Ca}^{2+}$ increase attributed to<br>Malpighian tubules (%) |
| --- | --- | --- | --- |
| 5 | 39.5 ± 13.5 | 14.7 ± 3.9 | 37.1 ± 10.0 |
| 10 | 44.6 ± 9.3 | 31.0 ± 3.2 | 69.6 ± 7.2 |
| 20 | 53.3 ± 17.3 | 35.0 ± 5.2 | 65.7 ± 9.8 |
| 40 | 46.5 ± 13.5 | 27.6 ± 6.9 | 59.4 ± 14.9 |
| 80 | 47.6 ± 16.6 | 22.9 ± 4.85 | 48.1 ± 10.2 |

**Table S3.** Estimated total amount of Ca^2+^ ingested per mosquito after 6 days of feeding on elevated dietary Ca^2+^ in 10% sucrose compared to the increase of Ca^2+^ in mosquito whole bodies (Supplemental Table 2). Amount of Ca^2+^ ingested was calculated by the product of Ca^2+^ concentration and volume ingested. The difference between the estimated ingestion and whole body increase is hypothesized to be excreted. Values are means ± s.e.m.

| $\text{CaCl}_2$<br>Concentration<br>(mM) in 10%<br>sucrose | Volume of<br>10% sucrose<br>ingested over<br>6 days<br>($\mu\text{L}/\text{mosquito}$) | Amount of $\text{Ca}^{2+}$<br>ingested over 6<br>days<br>(nmol/mosquito) | Whole body $\text{Ca}^{2+}$<br>increase<br>(nmol/mosquito) | Hypothesized<br>$\text{Ca}^{2+}$ Excreted<br>(nmol/mosquito) |
| --- | --- | --- | --- | --- |
| 5 | $11.4 \pm 1.2$<br>(n = 5) | $56.8 \pm 6.2$ | $39.5 \pm 13.5$ | $\approx 17.3$ |
| 10 | $13.1 \pm 0.6$<br>(n = 5) | $130.9 \pm 5.8$ | $44.6 \pm 9.3$ | $\approx 86.3$ |
| 20 | $7.8 \pm 0.4$<br>(n = 5) | $155.5 \pm 7.2$ | $53.3 \pm 17.3$ | $\approx 102.2$ |
| 40 | $6.1 \pm 0.8$<br>(n = 5) | $245.0 \pm 31.1$ | $46.5 \pm 13.5$ | $\approx 198.5$ |
| 80 | $7.5 \pm 2.3$<br>(n = 5) | $436.5 \pm 98.5$ | $47.6 \pm 16.6$ | $\approx 388.9$ |

**Figure S1.**
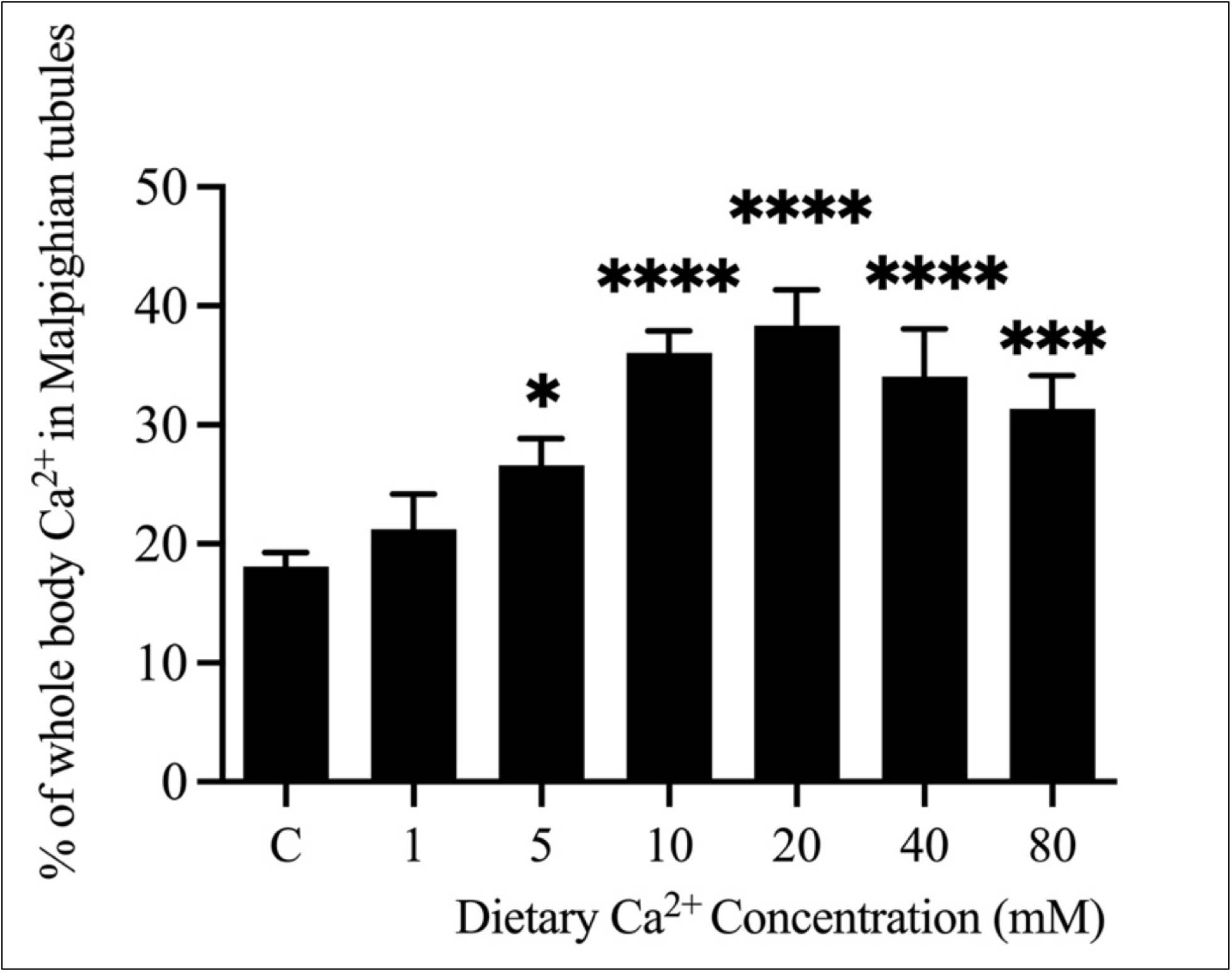
Percentage of whole body Ca^2+^ levels detected in Malpighian tubules. C = control group and numbers indicate Ca^2+^ concentration (mM) in treated groups. Percentages were determined by dividing the tubule Ca^2+^ content for each biological replicate by the respective mean whole body Ca^2+^ content within a group. Values are mean ± s.e.m; n = 16, 5, 7, 7, 8, 7, and 6 respectively, for C, 1, 5, 10, 20, 40, and 80 mM Ca^2+^ treatments

**Figure S2.**
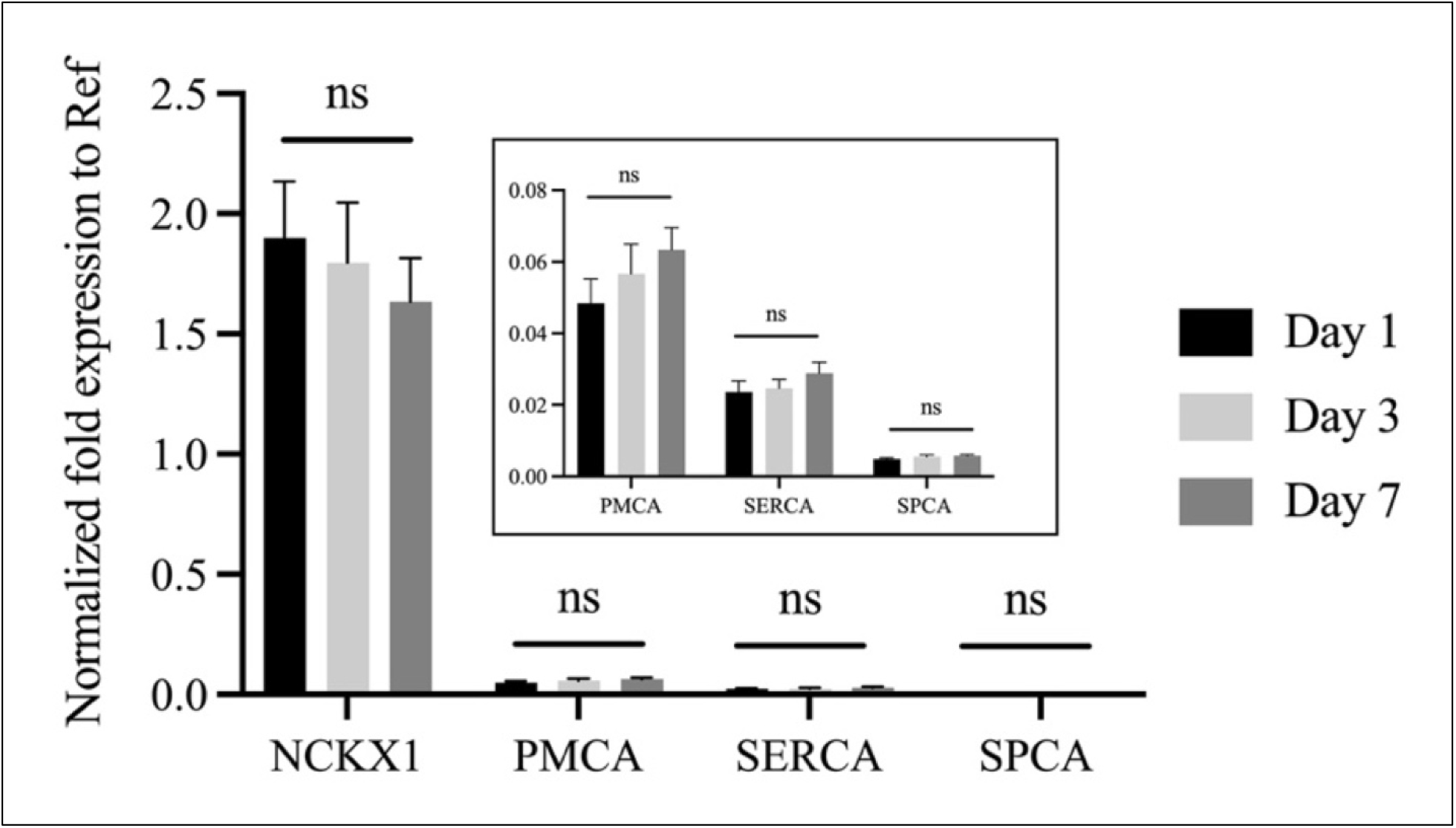
Relative mRNA expression of Ca^2+^-transporters in Malpighian tubules of controls; inset shows magnified view for *PMCA*, *SERCA*, and *SPCA*. Black, light grey, and dark grey bars respectively represent the relative expression values of control groups at 1 d, 3 d, and 7 d after feeding on 10% sucrose diets. PMCA: p = 0.1553 (Kruskal Test); SERCA, SPCA, and NCKX: p = 0.4294, 0.2492, 0.7074 (Ordinary One-way ANOVA). Values are mean ± s.e.m; n = 5 replicates.

